# Open EGGbox: an open-source 3D-printed embryonic *Gallus gallus* toolbox for electroporation and culture/live imaging of avian embryos *ex ovo*

**DOI:** 10.1101/2025.03.04.641180

**Authors:** Melissa Antoniou-Kourounioti, Felícitas Ramírez de Acuña, Raphael Caspar Constantin Schettler, Sakshi Kaushal Udar, Aitor Hamzic Petite, Ramayee Vidhya Sivasubramanian, Andrea Erika Münsterberg, Timothy Grocott

## Abstract

We present an open source, 3D-printed toolbox for avian embryology. The toolbox includes an electroporation chamber for transfecting functional molecular reagents into developing embryos, and a set of live-imaging chambers, which support avian embryo development while presenting them to a wide variety of microscope setups. We demonstrate both electroporation and imaging chambers by transfecting novel fluorescent reporter constructs for the TGF-beta signalling pathway and Pax7, Brachyury, Cdx2 and Sox2:Oct4 transcription factors into chick embryos and performing time-lapse imaging from stages HH3 - HH12 via both widefield and confocal fluorescence microscopy. Open-source code and ready-to-print STL files are freely available from a GitHub repository in line with FAIR (findability, accessibility, interoperability, and reusability) principles.

## Introduction

The ability to culture and manipulate avian embryos *ex ovo*, such as those of the chick *Gallus gallus*, affords unsurpassed access for investigating development within amniota (reptiles, birds, mammals). A variety of bespoke technical approaches and apparatus have grown up around the chick model. The motivation behind this study was to consolidate some of the most generally useful approaches into an open and standardised toolkit for the avian embryology community, reducing the effort needed to adopt the chick as an experimental model.

A number of *ex ovo* culture methods have been developed for chick, culminating in Dennis New’s method of supporting the vitelline membrane by means of a metal or glass ring (New, 1955; Stern and Bachvarova, 1997). This approach was later streamlined in the form of Early Chick (EC) culture, wherein the metal/glass ring was substituted with a disposable filter paper ring, simplifying the process of preparing embryos for culture (Chapman et al., 2001; Streit, 2009). EC culture is perhaps the most widely used method for observing early avian development via time-lapse microscopy. A complimentary *ex ovo* culture approach is the Cornish pasty method (Connolly et al., 1995; Nagai et al., 2011), which forgoes the vitelline membrane entirely, permitting development to later stages, but making live-imaging difficult due to the unstable nature of the free-floating embryo.

Once explanted, avian embryos are not only accessible for observation, but also for a wide variety of functional perturbations including microsurgery/grafting, bead implantation, and transient transfection/transgenesis via electroporation-mediated gene transfer (Mok et al., 2015; Streit et al., 2012). While these techniques are also available at later stages *in ovo*, their application to fragile earlier stages is far more tractable *ex ovo*, and this is particularly true for whole-embryo electroporation.

### Electroporation

Electroporation-mediated gene transfer is a widely used method for performing genetic gain- and loss-of-function experiments in avian embryos including the chick. With a suitable electrode setup, this technique may be performed either *in ovo* or with explanted embryos cultured *ex ovo* and is commonly used in combination with the EC culture method. Early-stage embryos isolated and electroporated with the EC method may subsequently be transferred into Cornish Pasty culture for longer-term culture.

*Ex ovo* electroporation may be combined with other functional techniques including tissue grafting/ablation, bead implantation, explant cultures, cell dissociation/sorting for genomic/transcriptomic analyses, fate mapping and time-lapse imaging. Although mesenchymal tissues are more difficult to electroporate than epithelia such as ectoderm, their progenitors may be efficiently electroporated prior to undergoing epithelial-mesenchymal transition whilst still resident within the epithelial epiblast. The practised experimentalist can use fate maps to target particular cell lineages within the epiblast, but transgene expression may also be targeted and timed via carefully chosen or designed *cis*-regulatory elements.

Many labs that practise electroporation employ hand-made electrodes or expensive commercial alternatives. Often these are simple platinum or other wires (Voiculescu et al., 2008), but uniform electroporation targeting large areas of the epiblast requires parallel plate electrodes (Uchikawa et al., 2003; Uchikawa et al., 2017; Williams and Sauka-Spengler, 2021). While some commercial apparatus exists, expense and availability can be prohibitive.

### Live imaging of EC-cultured embryos

The EC culture method (Chapman et al., 2001; Streit, 2009) is often used to present live avian embryos to a microscope objective for live time-lapse imaging (Cui et al., 2005; Song et al., 2011) including the submerged filter paper sandwich variation (Schmitz et al., 2016). EC-cultured embryos may be incubated in multi-well plates enabling several embryos to be imaged at a time (Song et al., 2011). A challenge with this approach is a conflict between the depth of substrate (albumin or albumin/agar) required for healthy development versus objective working distance (i.e. distance between the lens surface and the focal point). Conventional plastic-bottom multi-well plates are much thicker than a glass coverslip and offer poorer optical properties. This limits the potential for high-resolution imaging using large numerical aperture/short-working distance objectives. Single-use glass-bottomed dishes are available, but while suitable for inverted microscopes, the experimentalist may require an upright instrument, e.g. to take advantage of high numerical aperture dipping objectives.

The challenge of presenting *ex ovo* cultured chick embryos to upright objectives was previously tackled via fabrication of single-use imaging chambers (Voiculescu and Stern, 2012) but as with electroporation chambers, their fabrication is complex, time-consuming, and presents barriers to reproducibility and wider adoption. The related filter paper sandwich approach permits single embryos to be cultured fully-submerged (Schmitz et al., 2016) without need for complex apparatus, but can be difficult to extend to multi-embryo experiments.

### Open EGGbox

Here we present an open-source and FAIR (findable, accessible, interoperable, reproducible) 3D-printed electroporation chamber, EGGbox-E, intended as a standardised apparatus to aid reproducibility and widen access to this powerful technique. This chamber was designed to be compatible with existing detailed methodologies for *ex ovo* electroporation of chick embryos (Uchikawa et al., 2017; Voiculescu et al., 2008; Williams and Sauka-Spengler, 2021; Yukinori, 2012).

We additionally present a series of open-source and FAIR 3D-printed re-usable culture/imaging chambers that can support one or more chick embryos in EC and filter paper sandwich culture, while presenting them to microscope objectives in both upright and inverted configurations.

Open EGGbox comprises both the physical apparatus and the open-source version-controlled code used to create them. This may be freely accessed, modified and redistributed under an open-source licence.

### Synthetic reporter genes

We demonstrate Open EGGbox by electroporating chick embryos *ex ovo* with an array of novel synthetic reporter constructs designed to drive fluorescent reporter gene expression in different cell lineages or else report specific signalling pathway activities. We show that Open EGGbox can facilitate time-lapse imaging of broad, targeted and lineage-specific transgene expression via both widefield and confocal imaging in live chick embryos.

## Results

### 3D-printed electroporation chamber: EGGbox-E

#### Design & Assembly

The electroporation chamber (Fig. 1A, B) was adapted from a design by Tatjana Sauka-Spengler (Williams and Sauka-Spengler, 2021) to permit 3D-printing, easy assembly/disassembly, and increased volume to accommodate larger filter-paper EC culture rings or glass New culture rings. Larger rings provide more room for epiboly during culture, prolonging the window during which normal morphogenesis may be studied.

**Figure 1:**
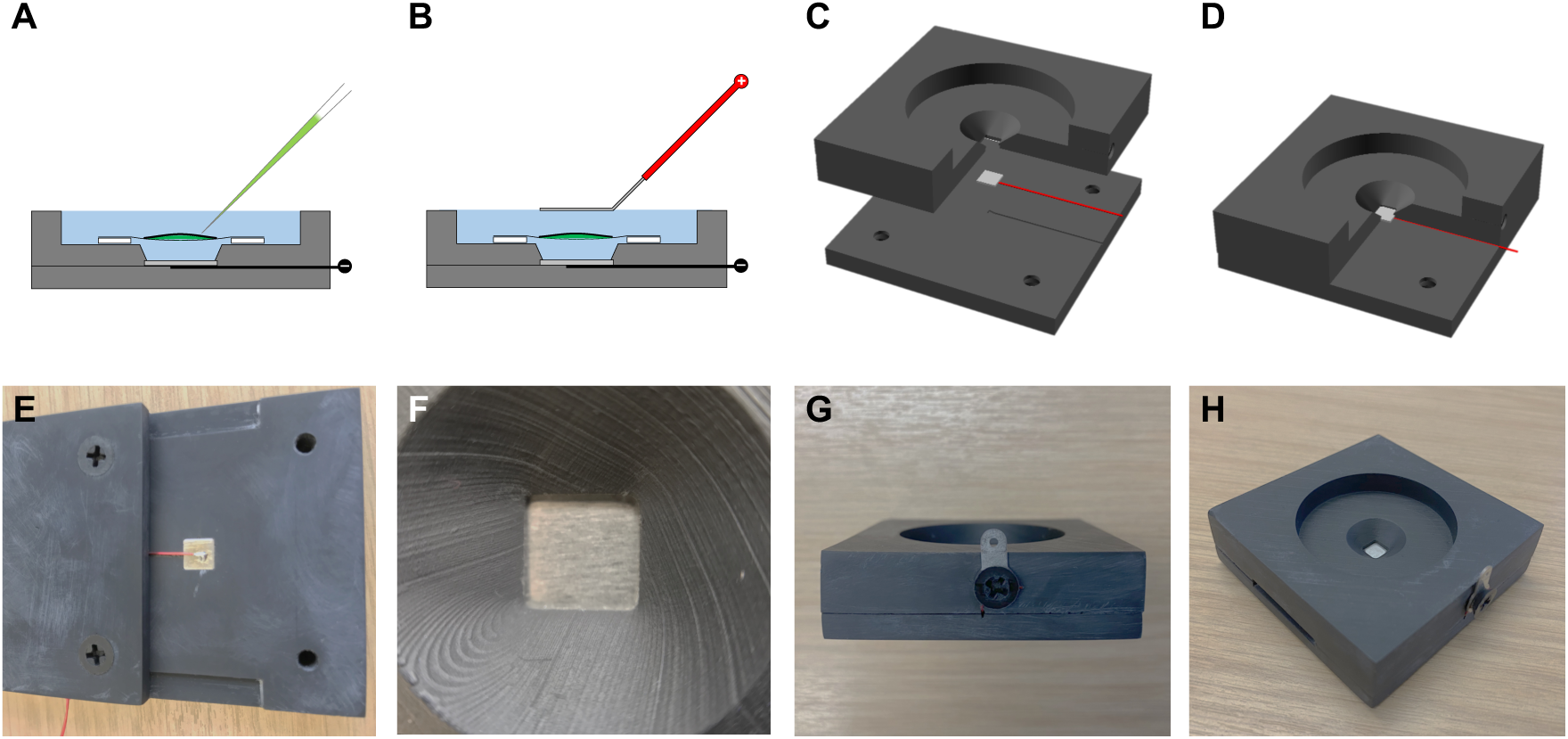
EGGbox-E 3D-printed electroporation chamber. **A-B)** Conceptual schematic of the electroporation chamber. Not to scale. **A)** Embryo supported by filter-paper ring is positioned ventral side up, over the negative plate electrode inside the electrolyte-filled chamber. DNA etc. (green) is microinjected into the space between the vitelline membrane and embryo. **B)** Positive paddle electrode is positioned above the embryo and the DNA is electroporated upwards into the dorsal side (epiblast). **C-D)** 3D models of the electroporation chamber. Cutaways show interior detail. The plate electrode is recessed into the upper half, while the attached wire is recessed into the lower half and exits at one side. **C)** Exploded view. **D)** Assembled view. **E-H)** Assembly of the 3D-printed chamber. **E)** Fitting the plate electrode. The platinum plate is cut slightly oversized, then filed down to exactly fit the recess in the underside of the upper half. The lower half is fitted off-centre to hold the connecting wire in place while soldering to the plate. **F)** Top view: platinum plate electrode after fitting. **G)** Side view: connecting wire exits the side of the chamber, where it connects to a terminal screwed to the chamber’s side. **H)** The assembled chamber.

The chamber comprises upper and lower halves, between which a cathode electrode is sandwiched (Fig. 1C, D). The cathode itself is fashioned from platinum foil (5 x 5 x 0.5 mm) to which a narrow connecting wire (0.4 mm diameter) is soldered (Fig. 1E, F). The completed assembly is screwed together and may be dismantled for deep cleaning. See methods for further details.

We explored different 3D printing processes and materials before settling on high-resolution SLA resin. Some examples were printed using high temperature and dental resins, which may be autoclaved. However, we found the post-print UV curing process required for thermal stability led to undesirable warping of the finished parts. Thus, we settled on a standard high-detail resin, with a dark grey colour providing excellent contrast for observing embryo morphology under reflected illumination.

In addition to the 3D-printed chamber, a separate anode electrode is required. The anode may be fabricated from platinum plate similar to the cathode, but it is simpler to crush thick platinum wire (0.8 mm diameter) in a clamp to form a flattened end which can be bent at a 45-degree angle. See methods and (Williams and Sauka-Spengler, 2021) for further details.

#### Use

Once the chamber is assembled (Fig. 1G, H) both anode (paddle electrode) and cathode (chamber electrode) are connected to an electroporation unit (anode to positive, cathode to negative forming an electrolytic cell). The chamber’s upper half forms a well, which is filled with an electrolyte solution (e.g. Pannet-Compton, Ringer’s or Tyrode’s saline) and into which the explanted embryo is placed, ventral side up (Fig. 1A). Reagents to be transfected (e.g. plasmid DNA, antisense morpholino oligonucleotides; green in Fig. 1A) are micro-injected between the epiblast and vitelline membrane. The anode is then placed at the surface of the saline and negatively charged reagents are electroporated upwards towards the anode and into the dorsal side of the epiblast (Fig. 1B).

In use, the chamber is filled to the brim with electrolyte solution and the upper paddle electrode just contacts the surface to ensure a consistent inter-electrode distance yielding good reproducibility between embryos/experiments (Fig. 1B). For uniform targeting of broad areas, the upper electrode should be held parallel to the lower electrode to ensure a uniform voltage gradient (and thus transfection efficiency) across the embryo. When used in conjunction with Intracell OvoDyne electroporation units, we find the following electroporation settings produce consistent and efficient transfection (of both DNA constructs and morpholinos) and good embryo survival with primitive streak stage HH3/4 embryos: 10 volts (voltage gradient = 1 volt/mm), 5 pulses, 50 ms length, 100 ms space. We recommend experimenting with lowering/raising the voltage gradient slightly for younger/older embryo stages, respectively.

Once electroporated, embryos may be transferred to one of the EGGbox culture/imaging chambers/plates for live-imaging (see below), cultured in a 35 mm dish or 6-well plate on albumin (Williams and Sauka-Spengler, 2021) or albumin/agar (Chapman et al., 2001), or detached from their vitelline membranes by submerging them in a Ca2+/Mg2+-free saline (e.g. Ca2+/Mg2+-free Tyrode’s) for use in Cornish Pasty culture (Connolly et al., 1995; Nagai et al., 2011).

### 3D-printed culture/imaging chambers: EGGbox-1, EGGbox-D1, EGGbox-D6, & EGGbox-8

#### Design

Building on previous single-use designs intended for New’s culture (Stern, 1990; Voiculescu and Stern, 2012), we set out to develop a re-usable culture/imaging chamber that could support a single embryo in EC culture on both upright and inverted microscope stands (Fig. 2A, B). The 2-part design included i) a main chamber piece carrying a coverslip, a thin layer of albumin-agar (or just albumin), and a filter-paper ring bearing the embryo; ii) a translucent screwcap (to permit transmitted illumination) with vent holes to prevent pressure build-up popping the coverslip. The universal screwcap fits into the main application-specific chamber piece to secure the filter paper ring/embryo regardless of orientation. The screwcap is designed to carry a small volume of humidifying liquid in both upright and inverted orientations to prevent dehydration. The ability to culture embryos in upright configuration permits a variation of the application-specific chamber piece that includes an outer reservoir suitable for use with dipping objectives (Fig. 2C). Figure 2 D-F shows the corresponding 3D-printable designs (cutaways show internal details).

**Figure 2:**
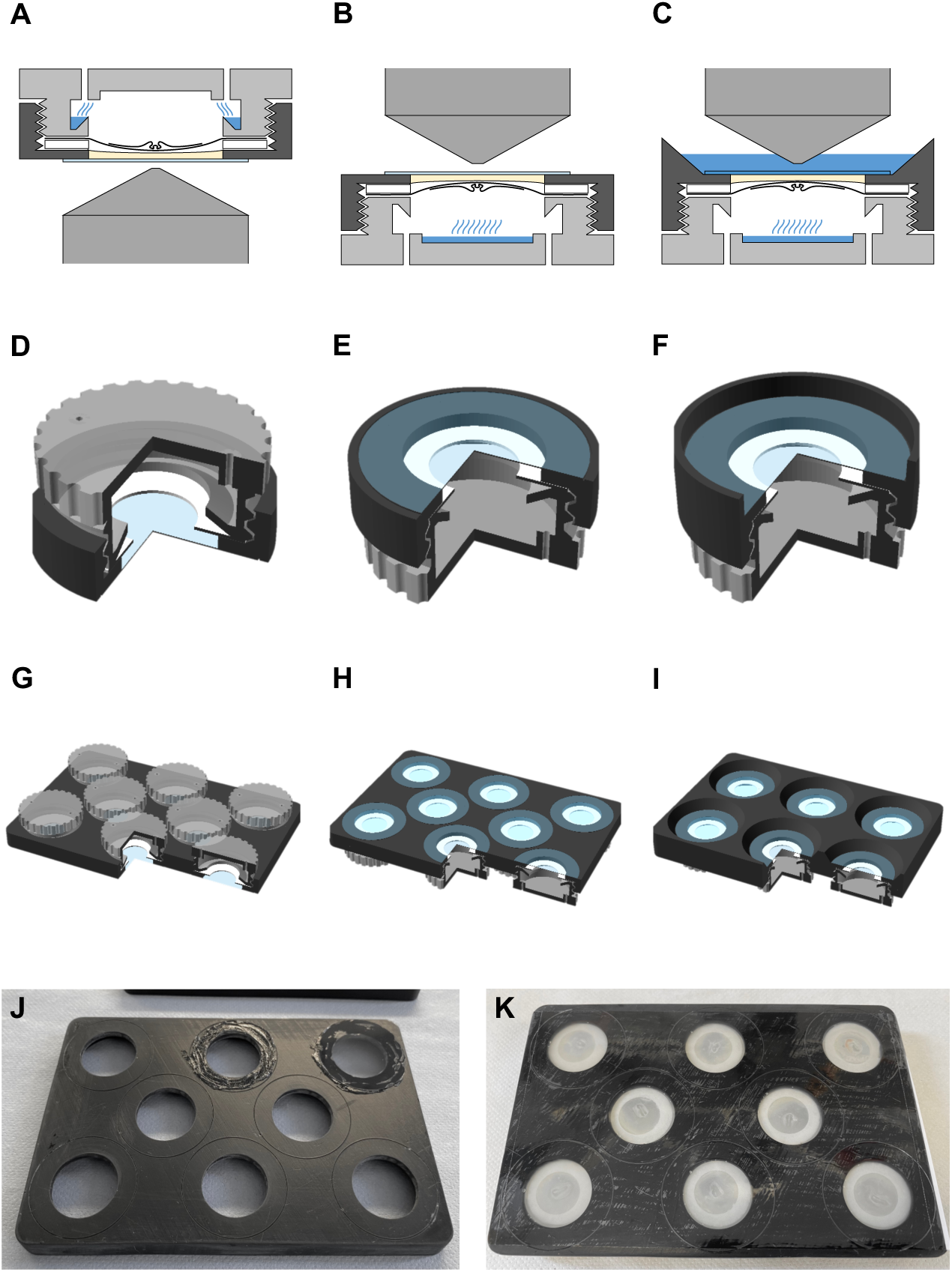
EGGbox 3D-printed culture/imaging chambers and plates. **A-C)** Concept for a 3D-printable multi-configuration culture/imaging chamber. Not to scale. An application-specific “lid” (dark grey) is fitted with a coverslip and filled with the preferred growth substrate (pale yellow), while a screw-in universal base (light grey) forms a humidified chamber. A filter paper ring bearing an embryo is sandwiched between the two, enabling incubation in upright or inverted configuration. **A**) Inverted configuration. **B)** Upright configuration. **C)** Upright configuration with “lid” designed for dipping objectives. **D-F)** 3D-printable single-embryo culture/imaging chambers. Cutaways show interior detail. **D**) EGGbox-1, inverted configuration. **E)** EGGbox-1, upright configuration. **F)** EGGbox-D1 for dipping objectives, upright configuration. **G-I)** 3D-printable multi-embryo culture/imaging plates. Cutaways show interior detail. **G-H)** EGGbox-8, 8-embryo culture/imaging plate, **G)** inverted configuration, **H)** upright configuration. **I)** EGGbox-D6, 6-embryo culture/imaging plate for dipping objectives, upright configuration. **J)** Coverslips are fixed to the outer surface of EGGbox-8 using vacuum grease before the growth substrate, embryos and screw caps are added from the other side. **K)** Imaging-quality self-adhesive plastic film may be used with EGGbox-8 instead of coverslips.

Economies of scale can be leveraged if multiple embryos are imaged simultaneously. We therefore extended our designs to a mutliwell plate format to increase the number of embryos that can be imaged per experiment. EGGbox-8 (Fig. 2G, H) provides 8 separate chambers in which embryos may be cultured upright or inverted, while EGGbox-D6 (Fig. 2I) enables culture/imaging of up to 6 embryos with a dipping objective on an upright stand. Multiple separate chambers help to isolate embryos both optically and in terms of potential microbial contamination, while also minimising dehydration. The internal dimensions of all chamber designs are identical as are the universal screwcaps, satisfying the requirement for interoperability.

#### Assembly & Use

Circular coverslips are reversibly attached using high vacuum grease (Fig. 2J). Alternatively, individual coverslips may be substituted by a single imaging-quality self-adhesive film (Fig. 2K). Coverslips provide superior sample stability and optical properties, whereas self-adhesive film is less messy and faster to setup/cleanup but affords a lower degree of sample stability in the z axis. Maintaining the focal plane as the film flexes during long-term imaging can be achieved via software autofocus (used here), or a hardware-based solution such as Definite Focus, although the latter doesn’t account for embryo movement relative to the coverslip/film.

Once coverslips/film are in place, the plates are inverted and wells are filled with albumin or albumin-agar from the inside, followed by the filter paper rings bearing vitelline membranes and attached embryos. The volume of semi-solid albumin-agar determines whether the substrate surface is concave, flat, or convex (e.g. 250 µl, 500 µl, or 750 µl, respectively), and consequently, the embryos’ eventual distance from the objective. A convex profile helps excess liquid to drain away from the embryos, facilitating attachment or re-attachment to the vitelline membrane as in New’s culture (New, 1955). This permits grafting of other avian embryos (e.g. quail) onto chick vitelline membranes that are loaded into the chamber (Stern and Bachvarova, 1997) in addition to homo-/hetero-chronic/topic tissue grafting between donor and host embryos/species. The trade-off is that increased substrate volume requires longer working-distance objectives (certainly > 2 mm with 750 µl of albumin-agar), which generally have lower numerical apertures (i.e. lower resolution).

### Design of synthetic reporter genes for monitoring signalling pathway and transcription factor activities

We tested EGGbox electroporation and culture/imaging chambers in a variety of experimental configurations, targeting both broad and localised areas of the chick epiblast at different stages, and a variety of different cell lineages using ubiquitous and lineage-specific reporter constructs. To this end, we developed a series of novel fluorescent reporter constructs from well-described transcription factor binding motifs (i.e. not derived from natural *cis*-regulatory elements).

The strategy for engineering these reporters is shown in Supplementary Figure 1 (Supp Fig. 1A-C), together with a summary of all reporters generated for this study (Supp Fig. 1D). Briefly, dsDNA oligos bearing specific binding motifs and pseudo-random spacers (Supp Fig. 1A) are assembled into fluorescent reporter plasmids via Golden Gate assembly (Supp Fig. 1B). Iterative amplification steps (Supp Fig. 1C) can exponentially grow the number of motifs to combat saturation of the transcription factor binding motifs and thus expand the reporters’ dynamic range and increase sensitivity to weaker transcriptional activators.

To validate this cloning strategy, we first re-created a destabilised canonical Wnt reporter by cloning 16 x LEF1 binding motifs upstream of a thymidine kinase promoter and driving expression of a destabilised Citrine fluorescent protein (called pTK-LEFx16-d1Citrine; Supp Fig. 1D, Fig. 3A). Destabilisation was achieved by fusing Citrine (Griesbeck et al., 2001) to a PEST-containing C-terminal fragment of mouse ornithine decarboxylase that reduces GFP half-life from ∼22 hours to ∼1 hour in mammalian cells (Li et al., 1998).

This reporter was co-transfected with an mCherry (Shaner et al., 2004) expression construct (pCICherry) (Grocott et al., 2020) as positive control into stage HH5 chick embryos *ex ovo* using the EGGbox-E electroporation chamber. Embryos were then transferred into EGGbox-8 culture/imaging chamber for culture. Widefield timelapse imaging was performed overnight using a Zeiss Observer 7 inverted microscope with environmental control.

As seen in Figure 3B-G, LEFx16-d1Citrine activity is observed in the posterior paraxial mesoderm exiting the primitive streak during stage HH6. This expression continues in the posterior paraxial (pre-somitic) mesoderm through to stage HH10-but was not observed in the forming somites (Fig. 3D-G).

**Figure 3:**
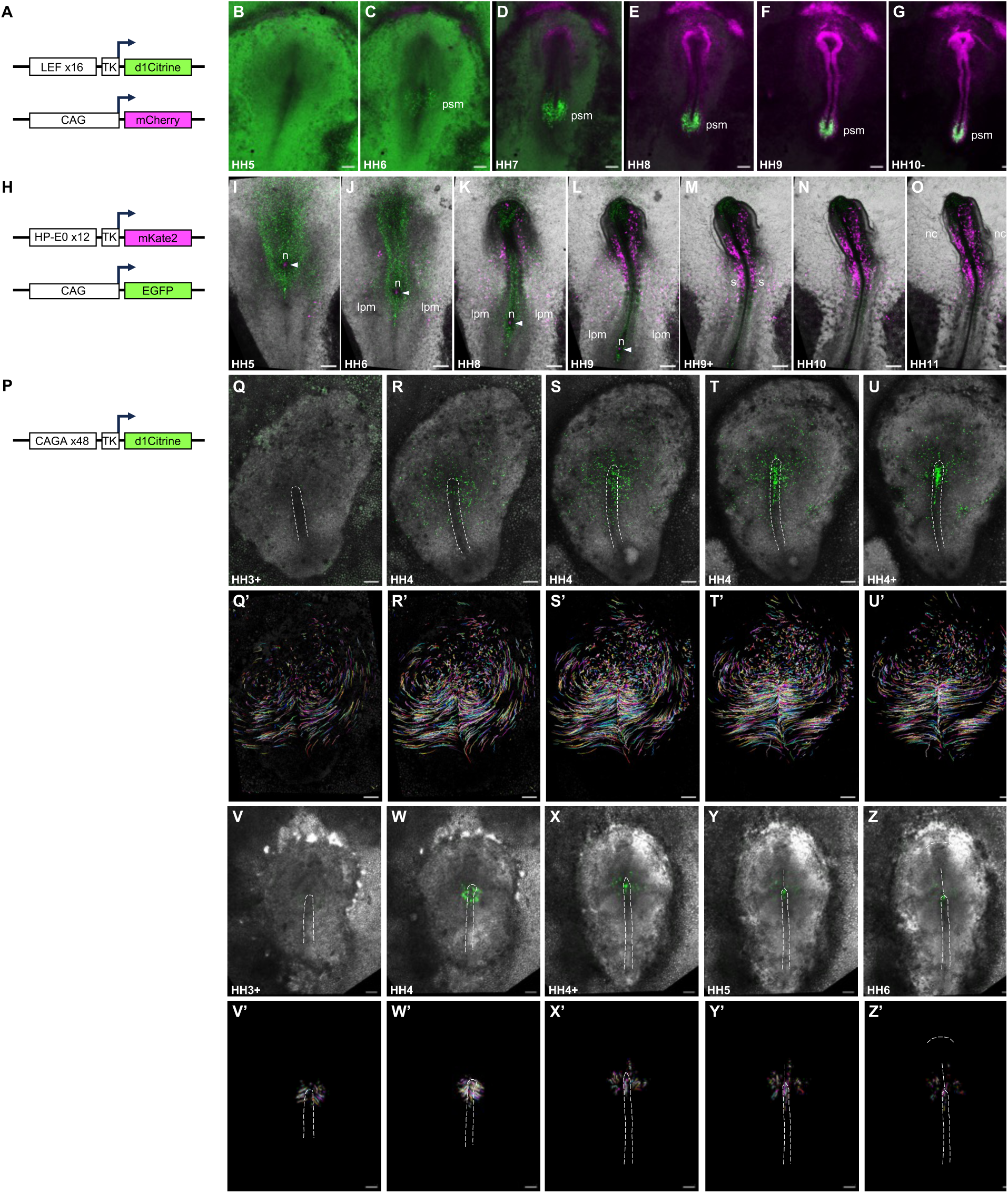
Electroporation and time-lapse imaging of synthetic reporters. **A)** A destabilised Wnt reporter comprising 16 x LEF1 binding motifs driving d1Citrine expression was co-electroporated with a CAG-mCherry targeting control using EGGbox-E and **B-G)** imaged on a Zeiss Observer 7 widefield microscope using EGGbox-8 between stages HH5 and HH10-. Scale bars = 500 microns. **H)** A Pax7 pioneer-factor reporter comprising 12 x HP-E0 pioneer motifs driving mKate2 was co-electroporated with a CAG-mCherry targeting control using EGGbox-E and **I-O)** imaged on a Zeiss LSM980 confocal microscope using EGGbox-8 between stages HH5 and HH11. Maximum projection of confocal fluorescence signals merged with transmitted brightfield illumination. Arrowheads in I-L indicate a small cluster of node-resident Pax7+ cells regressing with the node. psm = pre-somitic mesoderm; n = Hensen’s node; lpm = lateral plate mesoderm; s = somties; nc = neural crest cells. Scale bars = 250 microns. **P)** A destabilised TGF-beta reporter comprising 48 x “CAGA” Smad3/4 binding motifs driving d1Citrine expression was electroporated at stage HH3. **Q-U)** Broad electroporation using EGGbox-E and time-lapse imaging using EGGbox-8 and a Zeiss Observer 7 widefield microscope between stages HH3+ and HH4+. **Q’-U’)** Movements of TGF-beta responding cells were tracked over time. Broken lines in Q-U outline the primitive streak. Scale bars = 250 microns. **V-Z)** Localised electroporation using EGGbox-E and time-lapse imaging using EGGbox-8 and a Zeiss Observer 7 widefield microscope between stages HH3+ and HH6. **V’-Z’)** Movements of TGF-beta-responding cells were tracked over time. Broken lines in V-Z’ outline the primitive streak, notochord and head fold. Scale bars = 250 microns.

### Demonstration of EGGbox-E and EGGbox-8 via electroporation and time-lapse imaging of novel reporters

We next generated a novel reporter to capture the pioneer activity of transcription factor Pax7, which is active in a number of ectodermal and mesodermal tissues. Pax7 pioneer function is mediated by a motif termed HP-E0 (Supp Fig. 1D), which is recognised by both the paired domain and paired-type homeodomain of Pax7 (Pelletier et al., 2021). The spatiotemporal activity of this motif has not been assessed in embryos. We cloned 12 x HP-E0 pioneer motifs upstream of a thymidine kinase promoter driving mKate2 (Shcherbo et al., 2009) fluorescent protein expression (called pTKT2-HP-E0x12-mKate2; Supp Fig. 1D; Fig. 3H). This was broadly co-electroporated into the epiblast of stage HH4 chick embryos with an EGFP control construct (pCIG) using an EGGbox-E electroporation chamber then transferred to an EGGbox-8 culture/imaging chamber. In this instance embryos were observed on a Zeiss LSM980 inverted confocal microscope. 3D confocal + 2D brightfield timelapse imaging was performed between stages HH5 and HH11, a merged 2D maximum projection of which is shown in Figure 3I-O. As can be seen, Pax7 pioneer activity was observed in a small cluster of node-resident cells (Fig. 3I-O, arrowhead) and lateral plate mesoderm (Fig. 3I-L), anterior somites (Fig. 3L-O), and cephalic neural crest (Fig. 3K-O).

To monitor TGF-beta signalling activity, we cloned 48 x “CAGA” Smad3/4 motifs (Dennler et al., 1998) upstream of a thymidine kinase promoter and driving expression of a destabilised Citrine fluorescent protein (called pTK-CAGAx48-d1Citrine; Supp Fig. 1D, Fig. 3P). This was broadly electroporated using an EGGbox-E electroporation chamber in stage HH4 chick embryos, which were then transferred to an EGGbox-8 culture/imaging chamber. Widefield timelapse imaging was performed using a Zeiss Observer 7 inverted microscope. As can be seen (Fig. 3Q-U), TGF-beta-responding cells were observed across the epiblast with the signal gradually increasing in intensity as they converged on the primitive streak. In tracking the movements of these TGF-beta-responding epiblast cells (Fig. 3Q’-U’), we observed the characteristic counter-rotational “polonaise” movements (Fig. 3Q’-S’), which became less evident as stage HH4 proceeded (Fig. 3T’-U’). In the latter half of gastrulation, there was negligible movement of TGF-beta-responding epiblast cells anterior of the primitive streak in the region of the prospective neural plate (Fig. 3T’-U’; fewer cell tracks, and short in length), whereas those TGF-beta-responding epiblast cells converging on the primitive streak were moving much further and faster in comparison (Fig. 3T’-U’; many long tracks).

To gain a clearer picture of TGF-beta signalling as cells pass through the primitive streak, we performed a more localised electroporation using EGGbox-E. Injection of the pTK-CAGAx48-d1Citrine TGF-beta reporter was targeted to a small region of epiblast centred on the anterior-most primitive streak. Embryos were then transferred to an EGGbox-8 culture/imaging chamber prepared with self-adhesive plastic film in place of coverslips (Fig. 2K). Time-lapse widefield imaging was performed on an inverted Zeiss LSM980 confocal instrument by opening the pinhole to maximum and using software auto-focus to compensate for any flexing of the plastic film.

As can be seen (Fig. 3V-Z’), the TGF-beta reporter activated shortly after electroporation in epiblast cells immediately surrounding the anterior primitive streak (Fig. 3V, V’; stage HH3+). As these cells converged towards the midline, there was a marked up-regulation of the fluorescence signal (Fig. 3W, W’; stage HH4). By stage HH4+, cells that passed through the anterior primitive streak (undergoing EMT) continued to show reporter activity, although fluorescence intensity appeared somewhat diminished (Fig. 3X, X’; stage HH4+). Whereas some TGF-beta-responsive cells remained within the primitive streak, those that passed through radiated out from the midline with reporter activity diminishing still further (Fig. 3Y, Y’; stage HH5). TGF-beta-responding cells that entered the anterior primitive streak at stage HH4 and exited during stage HH4+/5 ultimately contributed to paraxial mesoderm (Fig. 3Z, Z’; stage HH6).

To demonstrate broad targeting and confocal timelapse imaging of multiple cell lineages across a single embryo, we generated synthetic reporters driving expression of orthogonal fluorescent proteins in response to different transcription factors: Sox2:Oct4 (targeting pluripotent progenitors) with a destabilised Cerulean (Loh et al., 2006; Rizzo et al., 2004), Brachyury (targeting embryonic mesoderm) with mKate2 (Kispert and Herrmann, 1993; Shcherbo et al., 2009), and Cdx2 (targeting extraembryonic ectoderm and posterior mesoderm) with Citrine (Griesbeck et al., 2001; Huang et al., 2017) (Supp Fig. 1D; Fig. 4A). EGGbox-E was used to achieve broad electroporation of all three reporters into stage HH3 chick embryos, which were then transferred to an EGGbox-8 culture/imaging chamber prepared with glass coverslips. Imaging was performed from stage HH4 (Fig. 4B) onwards. As can be seen (Fig. 4C-H), Sox2:Oct4+ putative pluripotent progenitors (Fig. 4C-H, cyan) were evident in the anterior marginal zone at the junction of the areas pellucida and opaca at stage HH4 and spanning both areas pellucida and opaca either side of the marginal zone posteriorly. The number of Sox2:Oct4+ cells diminished as development proceeded to stage HH6. Conversely, Cdx2+ cells (Fig. 4C-H, yellow cells), while not evident at stage HH4, appeared in great numbers in the posterior area opaca. Meanwhile, Bra+ mesoderm progenitors (Fig. 4C-H, magenta) were observed converging on the posterior primitive streak throughout the movie.

**Figure 4:**
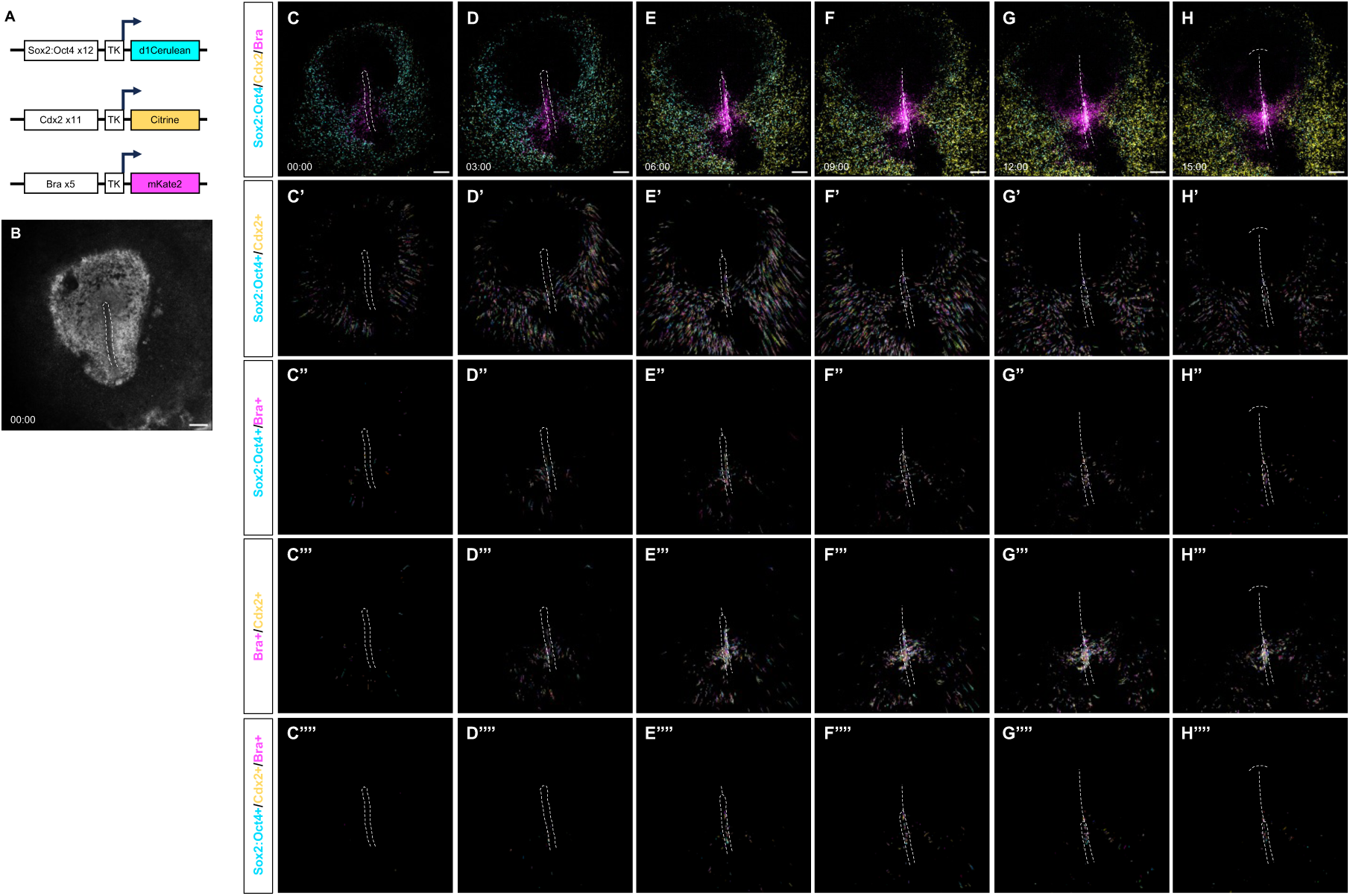
Co-electroporation and confocal time-lapse imaging of synthetic Sox2:Oct4, Brachyury and Cdx2 reporters. **A)** Synthetic enhancers with multiple copies of Brachyury, Cdx2, or Sox2:Oct4 composite binding motifs were coupled with different fluorescent reporter genes. **B)** All 3 reporters were co-electroporated into stage HH4 chick embryos using the EGGbox-E electroporation chamber. **C-H)** Time-lapse imaging using the EGGbox-8 culture/imaging plate on a Zeiss LSM980 confocal instrument. Cyan, Sox2:Oct4x12-d1Cerulean reporter; Yellow, Cdx2x11-Citrine reporter; Magenta, Brax5-mKate2 reporter. Broken lines indicate the locations of primitive streak, notochord and head fold (in order of appearance). Scale bars = 500 microns. **C’-H’’’’)** Trajectories of different sub-populations were tracked over time. **C’-H’)** Trajectories of Sox2:Oct4 and Cdx2 co-expressing cells. **C’’-H’’)** Trajectories of Sox2:Oct4 and Bra co-expressing cells. **C’’’-H’’’)** Trajectories of Bra and Cdx2 co-expressing cells. **C’’’’-H’’’’)** Trajectories of Sox2:Oct4, Bra and Cdx2 triple-positive cells.

We performed single cell tracking to gauge whether Sox2:Oct4+ progenitors could be observed transitioning into either Cdx2+ extraembryonic ectoderm or Bra+ embryonic mesoderm, and this was indeed the case (Fig. 4C’-H’’’’). Tracking of Sox2:Oct4+/Cdx2+ double-positive cells showed that the majority radiated out from the marginal zone to occupy the posterior area opaca, while a smaller number entered the posterior primitive streak (Fig. 4 C’-H’). Conversely, Sox2:Oct4+/Bra+ double-positive cells, which were fewer in number and slower-moving (shorter tracks), converged on the primitive streak (Fig. 4 C’’-E’’) before exiting laterally (Fig. 4F’’-G’’) and diminishing as Sox2:Oct4 expression was extinguished (Fig. 4H’’). A greater number of Cdx2+/Bra+ double-positive cells were observed within the primitive streak and exiting laterally (Fig. 4C’’’-H’’’), while a very few Sox2:Oct4+/Bra+/Cdx2+ triple-positive cells were mainly restricted to the streak itself (Fig. 4C’’’’-H’’’’). These observations suggest an expression sequence of Sox2:Oct4 > Bra > Cdx2 in posterior mesoderm progenitors as they first enter and then exit the streak.

As seen in Fig. 4D, we observed two distinct movements of Sox2:Oct4+ progenitors. The first of these was a centrifugal displacement of Sox2:Oct4+ cells situated in the marginal zone, corresponding to expansion of the blastoderm, some of which transitioned into Cdx2+ extraembryonic ectoderm (compare Fig. 4D, E). The second was a midline convergence and posterior displacement of Sox2:Oct4+ progenitors from the posterior marginal zone and into the primitive streak. These gastrulating progenitors were observed transitioning into either Bra+ mesoderm, or Bra+/Cdx2+ posterior mesoderm (compare Fig. 4D, E, F).

## Discussion

### The case for Open EGGbox

*Ex ovo* culture of avian embryos has contributed much to our understanding of amniote development, partly facilitated by a variety of methods for manipulating gene function in live embryos. These include mis-expression from plasmid expression vectors (e.g. constitutively actives, dominant negatives), mRNA knockdown via antisense RNAi, translation- or splice-blocking via morpholino oligonucleotides, introducing genetic lesions via CRISPR/Cas9, and modulation of endogenous gene function via CRISPR-ON/-OFF. In addition, engineered reporter genes have furnished avian embryologists with a means of monitoring cellular dynamics (e.g. FUCCI for cell cycle, Apoliner for apoptosis, etc.), cell lineages (e.g. Autobow for cell tracking), cell signalling (various pathway reporters), and gene regulation (via *cis*-regulatory elements, but also sensors engineered for specific *trans*-acting factors as shown here).

These powerful tools all require a means of introducing the requisite molecular reagents (plasmid vectors, morpholinos etc.) into cells of the developing embryo. By far the most popular method for transient *ex ovo* studies is electroporation-mediated gene transfer. In the case of reporter genes, there is also a need to monitor their activities in live embryos. While suitable apparatus and methods for *ex ovo* electroporation and live imaging of avian embryos have been reported previously, their development and adoption has been largely ad hoc. Here we consolidated previously successful methods and tools to develop and thoroughly test a standardised and open-source toolbox, Open EGGbox, for *ex ovo* electroporation and live imaging using the chick as a model organism (Fig. 5).

**Figure 5:**
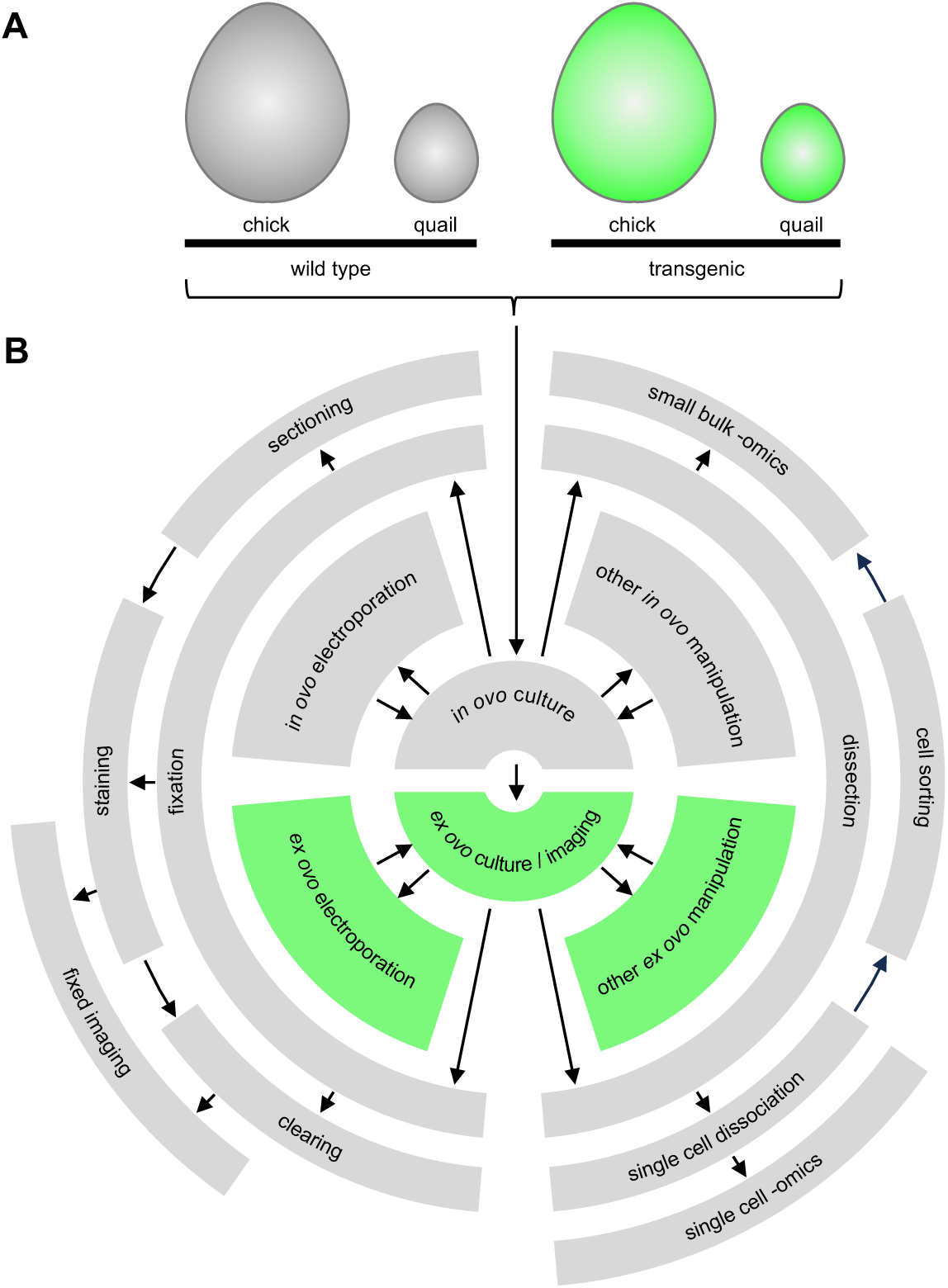
Application domain for Open EGGbox. **A)** In addition to wild type embryos, a growing number of fluorescent reporter lines are available for both chick and quail, **B)** both of which are amenable to a host of *in ovo* and *ex ovo* experimental approaches and downstream analyses. Arrows exemplify how various experimental methods may be sequentially applied to individual embryos. The *ex ovo* methods highlighted in B) correspond to the application domain for Open EGGbox.

Open EGGbox electroporation and culture/imaging chambers have been tested and routinely used in our lab over several years. When comparing the EGGbox-E electroporation chamber and the previously reported chamber on which the design was based (Williams and Sauka-Spengler, 2021), we have detected no difference in the proportion of embryos that survive and develop normally, nor the efficiency of electroporation. Likewise, embryo survival, rate of development and morphology was comparable between those cultured in the various EGGbox culture/imaging chambers versus conventional 6-well plates or individual 3.5 mm culture dishes. The materials used for 3D-prinitng have shown no signs of toxicity, although we ensured that 3D-printed parts were thoroughly washed prior to assembly and use. Nor have we observed any differences in embryo development when culture/imaging chambers were used in upright/inverted orientations or prepared with coverslips/plastic film. In sum, Open EGGbox has proven to be reliable toolkit for a variety of avian developmental studies.

### Complementarity with transgenic tools

Although developed with the chick in mind and hand, Open EGGbox may be used with other avian species, notably quail (Fig. 5A), which can thrive when grafted onto larger chick vitelline membranes (Stern and Bachvarova, 1997). Many transgenic reporter lines have been developed for both chick (Baek et al., 2018; Balic et al., 2014; Davey et al., 2018; Ho et al., 2019; McGrew et al., 2008; Oh et al., 2024; Rozbicki et al., 2015) and quail (Huss et al., 2015; Moreau et al., 2019; Saadaoui et al., 2020; Sato et al., 2010; Scott and Lois, 2005; Serralbo et al., 2020) (recently reviewed in (Henderson et al., 2025)) with an emphasis on live imaging. There is much scope for combining live imaging of these lines with electroporation-mediated functional perturbations, while Cre-inducible lines (Oh et al., 2024) may be electroporated with spatially and temporally specific Cre-drivers (Fig. 5B). Such drivers can be engineered similar to the novel fluorescent reporters presented here.

### Applications of synthetic reporters

We generated novel reporters to test electroporation and live imaging with Open EGGbox. These reporters (and their method of generation) will be generally useful for other developmental studies. For example, electroporate-able fluorescent reporter genes have been generated to monitor the activities of many developmentally important signalling pathways including Wnt, BMP, Hh, and Notch (Le Dréau et al., 2012; Rios et al., 2010; Toro-Tapia and Das, 2020; Vilas-Boas et al., 2011). However, to date none had been developed for monitoring the TGF-beta/Activin/Nodal pathway in live cells.

We were able to label different cell lineages by engineering synthetic reporters with motifs targeted by specific *trans*-acting factors – a kind of “painting by numbers”, where lineage-restricted activities of transcription factor motifs (the “numbers”) direct expression of different coloured fluorescent proteins (the “paints”). There is much scope for deriving novel transgenic lines that incorporate such reporters. However, these engineered reporters are most immediately useful as drivers for lineage-specific transgene expression in electroporation experiments.

### Limitations

Although the major components of Open EGGbox can be 3D-printed, some additional assembly is required. The most difficult aspect for many experimentalists will be assembling the electroporation chamber as this requires the cathode electrode to be fashioned and soldered. End users will also require a separate anode electrode. We have provided full parts lists (Supp Table 4, 5) and basic instructions (see Materials & Methods) for final assembly of both the electroporation chamber (cathode) and paddle electrode (anode). Where readers lack confidence in soldering, there are many good instructional videos online aimed at electronics hobbyists, and local engineering departments, workshops or hobbyists can likely help. Ultimately, it would be beneficial if assembly and use of these standardised apparatus were incorporated into established embryology training courses; indeed, Open EGGbox was recently demonstrated as part of the 2024 Edinburgh Gallus Genomics and Embryonic Development (EGGED 2024) Workshop alongside related technical approaches.

Many transcription factor binding motifs are shared by closely related family members. Thus, it is not necessarily the case that the intended *trans*-activator of an engineered reporter is the only *trans*-activator. However, it is possible to engineer more specific reporters using composite binding motifs that are bound cooperatively by two or more *trans*-acting factors. We demonstrated this with the Sox2:Oct4 reporter. Although Sox2 shares a core motif with other SoxB1 family members (some of which are expressed in the prospective neural plate at these same stages) and indeed Sox17 (a marker for definitive endoderm), our composite Sox2:Oct4 reporter instead showed only transient expression in the margin between the embryonic and extra-embryonic areas pellucida and opaca.

### Conclusion

A major intention of Open EGGbox is standardisation of both electroporation and live imaging methods, and thus enhanced reproducibility between experiments, individual lab members and the wider community. As much groundwork had been laid by other avian embryologists, a further intention was to broaden access and democratise these tools. The rapid proliferation of 3D-printing technologies in academic, commercial, and hobby spaces will help broaden access to these techniques, while the FAIR nature of the open-source code base will enable users to develop and share their own refinements or adaptations of these tools. One potential future development could be to incorporate fluidic channels into the EGGbox culture/imaging chambers to permit perfusion of morphogens or small molecules during live imaging.

## Materials & Methods

### Hen’s eggs

Fertile brown hen’s eggs (Henry Stewart) were incubated at 38°C in a humidified incubator until the required stage of development. The study was approved by the Animal Welfare & Ethical Review Board, School of Biological Sciences of the University of East Anglia, and all procedures were performed in accordance with the relevant guidelines and regulations.

### Reporter plasmid vectors

Novel reporter plasmid vectors were all derived from pTK-Citrine BsmBI LacZ v4 Nanotag_002 (Supp Fig. 2A), a kind gift from Tatajana Sauka-Spengler (Williams et al., 2019). pTK-d1Citrine (Supp Fig. 2B) was generated via Gibson Assembly (Gibson Assembly Master Mix, New England Biolabs), to insert an ODC degron at the C-terminus of the Citrine coding sequence. pTK-d1Cerulean (Supp Fig. 2C) was generated via Gibson Assembly to substitute the Citrine coding sequence of pTK-d1Citrine with that of Cerulean. pTKT2-mKate2 (Supp Fig. 2D) was generated via Gibson Assembly to i) substitute the Citrine coding sequence of pTK-Citrine with that of mKate2, and ii) flank the reporter cassette with left and right inverted terminal repeats (ITRs) to enable genome integration via Tol2 transposase. Supplementary Figure 2 shows plasmid maps for these reporter vectors, indicating the PCR primers/oligos used for Gibson Assembly, while Supplementary Table 1 lists the sequences of primer/oligo pairs.

### Construction of synthetic reporters

Synthetic enhancers were generated by first phosphorylating 5 µM each of combined sense/antisense ssDNA oligos (listed in Supp Table 2) using 0.5 units/µl T4 Polynucleotide Kinase (New England Biolabs) at 37 °C, then annealing via denaturation at 95 °C and gradual cooling to room temperature.

3.75 ng/µl of each annealed and phosphorylated dsDNA oligo was then combined with 3.75 ng/µl of reporter plasmid vector in a Golden Gate reaction containing 0.5 units/µl of BsmBI_v2 restriction endonuclease and 1 unit/µl of T4 DNA Ligase (New England Biolabs) with the following thermocycle: 42 °C 5 min, 16 °C 5 min, repeated for 60 cycles, 55 °C 5 min, 80 °C 20 min.

5 µl of the reaction volume was transformed into NEB 5-alpha chemically competent bacteria (New England Biolabs) using blue/white (X-gal) screening and antibiotic (carbenicillin) selection. White colonies were selected for miniprep, and synthetic enhancer sequences were verified via Sanger sequencing (Source Bioscience) in both directions using the primers pTK-Fwd (5’- AGGTACGGGAGGTACTTGG-3’) or pTK-Rev (5’-CAATGACAAGACGCTGGGC-3’).

### Computer aided design and 3D printing

All 3D printed parts are listed in Supplementary Table 3. All CAD work was performed and STL files generated using the free OpenSCAD software. STL files for all 3D-printed parts, together with OpenSCAD source code, are available from our GitHub site under a GPL3 licence: https://github.com/GrocottLab/Open-EGGbox

All 3D printing was performed using Formlabs SLA 3D printers and resins via a commercial printing service (Hexa-Cubed, Stamford, UK). Opaque parts were printed in Formlabs grey High Detail resin, providing a good contrast for visualising embryos in the electroporation chamber using reflected illumination. For the culture/imaging chambers, the opaque grey resin minimises photobleaching/phototoxicity of neighbouring samples on the same plate. The translucent universal screwcaps were printed in Formlabs clear High Detail resin to allow transmitted brightfield illumination during imaging. The softer egg rest in Supplementary Table 3 was printed in Formlabs silicone 40A resin. We explored printing both culture/imaging and electroporation chambers in Formlabs High Temperature and Dental resins, which may be autoclaved. However, the post-print UV curing process needed for these two materials resulted in some warping of the parts, and we found that autoclaving was not required in any case.

### Electroporation chamber assembly

The chamber was assembled upside-down. A 25 x 25 x 0.5 mm platinum/iridium foil (Pt80/Ir20) was cut down and filed into individual 5 x 5 x 0.5 mm plates and exactly fitted into a recess in the lower surface of the upper chamber half. The lower chamber half was screwed off-centre so that the wire channel in its upper surface could be used to clamp the enamelled copper wire in place. The connecting wire was flattened at the end and the enamel scraped off before clamping in place and soldering to the lower surface of the platinum plate using a temperature-controlled solder station (Fig. 1E). Flux was used to assist solder flow between wire and platinum plate. Once soldered, the lower half was refitted into its final position. The two-piece chamber was sealed by applying either a rubber sheet or silicon sealant beneath the platinum electrode. Enamel was removed from the copper wire that protruded from the chamber, and this was wrapped several times around a screw holding a terminal lug (Fig. 1G), which may be connected to the electroporation unit via a small crocodile clip. See Supplementary Table 4 for additional (non-3D-printable) parts.

### Paddle electrode assembly

The end 5 mm of a platinum wire (0.8 mm diameter x 20 mm long) was crushed in a clamp to form a flattened end that was then bent at a 45-degree angle. A connecting wire was passed through the bottom end of an empty pen barrel, soldered onto the non-flattened end of the platinum wire, and this assembly was then fed back into the bottom end of the barrel and glued in place with epoxy resin. A 4mm plug was attached to the unsoldered end of the connecting wire protruding from the top end of the pen barrel. See Supplementary Table 4 for the necessary parts and (Williams and Sauka-Spengler, 2021) for further details.

### Electroporation of chick embryos

The assembled chamber was connected to the negative terminal of an Intracel OvoDyne electroporation unit using a lead with small crocodile clip at one end and 4 mm plug at the other. The chamber well was filled to the brim with electrolyte solution (Ringer’s saline). Stage HH3/4 chick embryos prepared using the EC method were placed into the well ventral side up and centred over the chamber electrode. Plasmid DNA (∼2 µg/µl per plasmid) was microinjected through the blastoderm and into the space between dorsal epiblast and vitelline membrane using a glass capillary needle (Fig. 1A). To help visualise injection site/volume and control dispersal rate, DNA was combined with an electroporation buffer (0.1 % carboxymethylcellulose, 0.07 % Fast green, 1.5 mM MgCl_2_; used 1:10 for broad and 1:1 for localised injections) before injecting. A paddle electrode was connected to the electroporation unit’s positive terminal and positioned at the electrolyte surface directly above the embryo (Fig. 1B). Plasmid DNA was electroporated upwards into the epiblast layer using the following settings: 10 volts, 5 pulses, 50 ms width, 100 ms space. The electrolyte solution was replaced for each embryo.

Before each use, electrical polarity was checked by examining the connecting wires and electrical continuity was verified by performing a test electroporation without an embryo and watching for gas bubbles at cathode or anode. After each use, the chamber was thoroughly cleaned of residual egg yolk using fresh electrolyte solution and gently scraping all surfaces (especially corners) with a soft wooden toothpick. Once cleaned, the chamber was rinsed with distilled water then 70% ethanol and fully air-dried before the next use. For occasional deep cleaning the chamber was completely dismantled and the parts cleaned separately before reassembling. The paddle electrode was similarly cleaned after each use.

### Culture/imaging chamber assembly

High vacuum grease was carefully applied within the guidelines recessed into the outer surface of main chamber piece (Fig. 2J, top row centre) and a circular glass coverslip (Supp Table 5) was carefully pressed onto the grease (Fig. 2 top row right) ensuring a continuous seal and being careful that excess vacuum grease does not occlude the view through the coverslip. Alternatively, a single sheet on non-toxic low-autofluorescence plastic film (Supp Table 5) was applied to the entire outer surface (Fig. 2K).

The main chamber piece was inverted and each shallow well filled with 250 – 750 µl of albumin-agar prepared according to the EC method. An electroporated chick embryo was loaded into each well, ventral side up. Each well was sealed with a universal screwcap containing ∼250 µl of humidifying Ringer’s saline loaded into its inner edge (as in Fig. 2A). The chamber was maintained in this orientation for inverted imaging or carefully inverted for upright imaging.

After each use, the chambers were fully dismantled, the coverslips/film discarded, and the embryos discarded, fixed or cultured on as required. The chamber parts (plates and universal screwcaps) were thoroughly washed to remove residual vacuum grease and albumin/albumin-agar. Once washed, the parts were rinsed well with distilled water then 70% ethanol and fully air dried before the next use.

### Time-lapse imaging of chick embryos

EGGbox culture/imaging plates were imaged using Zeiss Observer 7 widefield and LSM980 + Airyscan2 confocal inverted microscopes with Zen Blue software. We defined a custom sample carrier to assist with sample location using the automated stage (Supp Fig. 3). Environmental chambers were set to 37.5 degC and images were captured at 5- or 10-minute intervals. The objectives used for this study were: EC Plan-Neofluar 2.5x 0.085 NA 8.8 mm working distance (Zeiss 420320-9902); Plan-Apochromat 10x 0.45 NA 2 mm working distance (Zeiss 420640-9900); EC Plan-Neofluar 20x 0.5 NA 2 mm working distance (Zeiss 423050-9900); EC Epiplan 20x 0.4 NA 3.2 mm working distance (Zeiss 422050-9961); EC Epiplan 40x 0.6 NA 2.2 mm working distance (Zeiss 422060-9901). For shorter working distance objectives, culture/imaging chambers were loaded with 250 µl of albumin-agar per well.

### Image analysis

All image processing and analysis was performed using ImageJ/Fiji (Schindelin et al., 2012). Timelapse images were registered to correct for embryo translation/rotation in xy using the plugin Linear Stack Alignment with SIFT Multichannel based on the method of (Lowe, 2004). Cell tracking was performed using the plugin TrackMate (Tinevez et al., 2017).

## Acknowledgments

We are grateful to Tatjana Sauka-Spengler and Octavian Voiculescu for helpful discussions and for sharing designs of their bespoke electroporation and imaging chambers. James Brydon of Hexa-Cubed gave valuable advice on printability of designs and materials choices. We thank James McColl and Paul Thomas of the Henry Wellcome Laboratory for Cell Imaging (UEA) for assistance with microscopy, and Victor Martinez-Heredia and Eirini Maniou for technical assistance. This work was funded by an Academy of Medical Sciences Springboard Award (SBF006\1154) to T.G., a Biotechnology and Biological Sciences Research Council (BBSRC) Responsive Mode grant (BB/N002970) to A.E.M., an Aid for the Re-qualification of the Spanish University System grant to F.R.d.A., a Gurdon/The Company of Biologists Summer Studentship to R.C.C.S, and a Genetics Society Summer Studentship to S.K.U.. We are grateful to the organisers and participants of the EMBO-funded EGGED 2024 workshop for the opportunity to demonstrate Open EGGbox and discuss future developments and applications.

## Author contributions

M.A-K. conceptualisation, methodology, software, validation, investigation, writing – review and editing. F.R.d.A. investigation, funding acquisition. R.C.C.S. investigation. S.K.U. investigation. A.H.P. investigation. R.V.S investigation. A.E.M. supervision, funding acquisition, writing – review and editing. T.G. conceptualisation, methodology, software, validation, formal analysis, writing – official draft preparation, writing – review and editing, visualisation, supervision, project administration, funding acquisition.

---

**Supplementary Figure 1:**
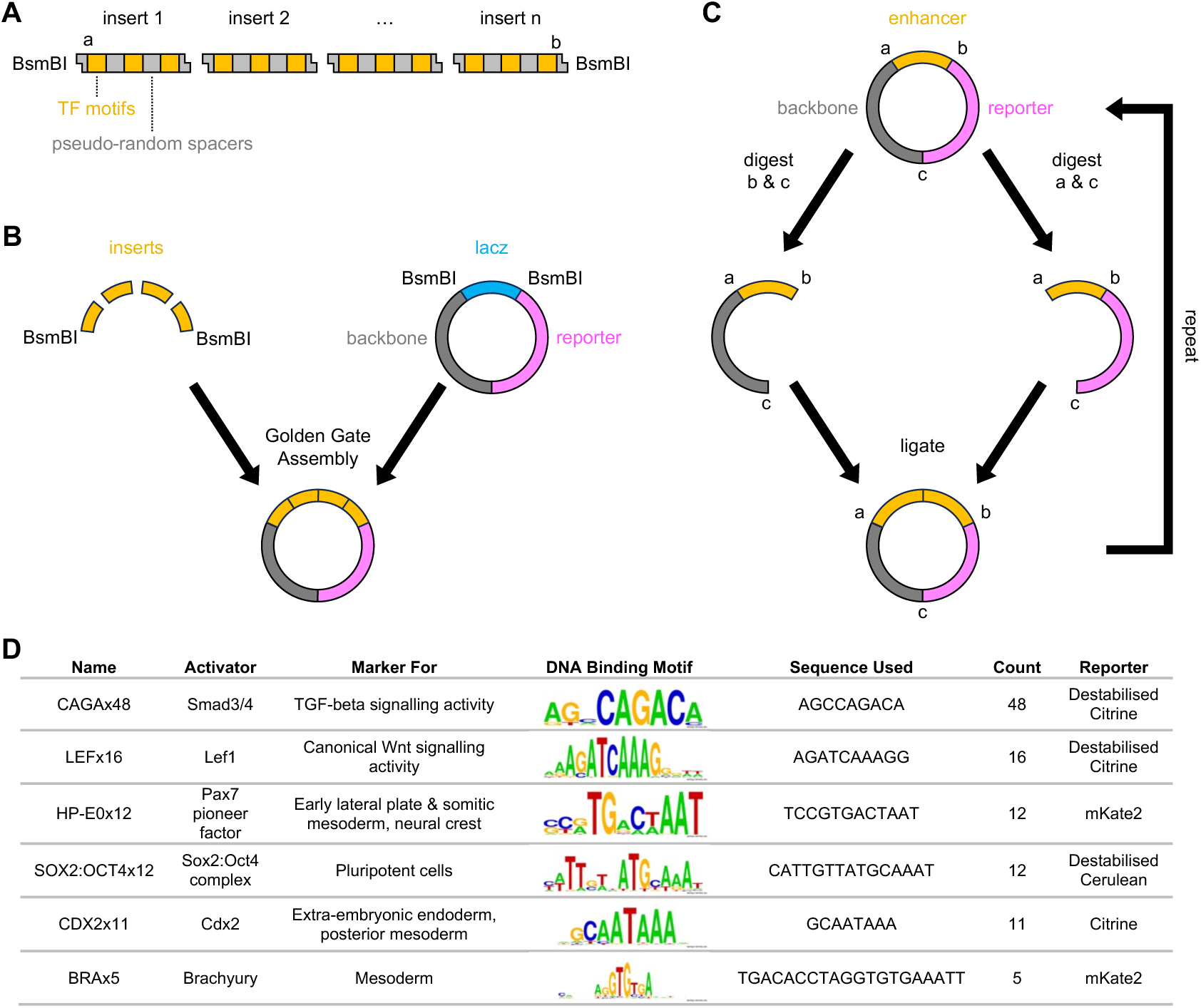
Construction of synthetic enhancers from known transcription factor (TF) binding motifs. **A)** One or more dsDNA inserts are assembled from annealed ssDNA oligos comprising multiple TF binding motifs separated by pseudo-random nucleotide spacers. To assist with Golden Gate Assembly, inserts incorporate uniquely complimentary 4 bp 5’ overhangs, the first and last overhangs being complimentary to BsmBI-generated overhangs within the target vector. To assist with later amplification, the first and last inserts also include distinct restriction enzyme sites (a & b) which generate complementary 5’ overhangs (e.g. AvrII, SpeI). **B)** The inserts are assembled with the target vector via Golden Gate Assembly using BsmBI. **C)** The number of TF motifs may be amplified by i) digesting the assembled plasmid with restriction enzymes a & c (e.g. AvrII & BamHI) or b & c (e.g. SpeI & BamHI), and ii) ligating the fragments bearing the synthetic reporter. Providing the composite a/b site cannot be re-cleaved, amplification may be repeated *ad infinitum*, doubling the number of TF motifs each round. **D)** Summary of synthetic reporters generated for this study.

**Supplementary Figure 2:**
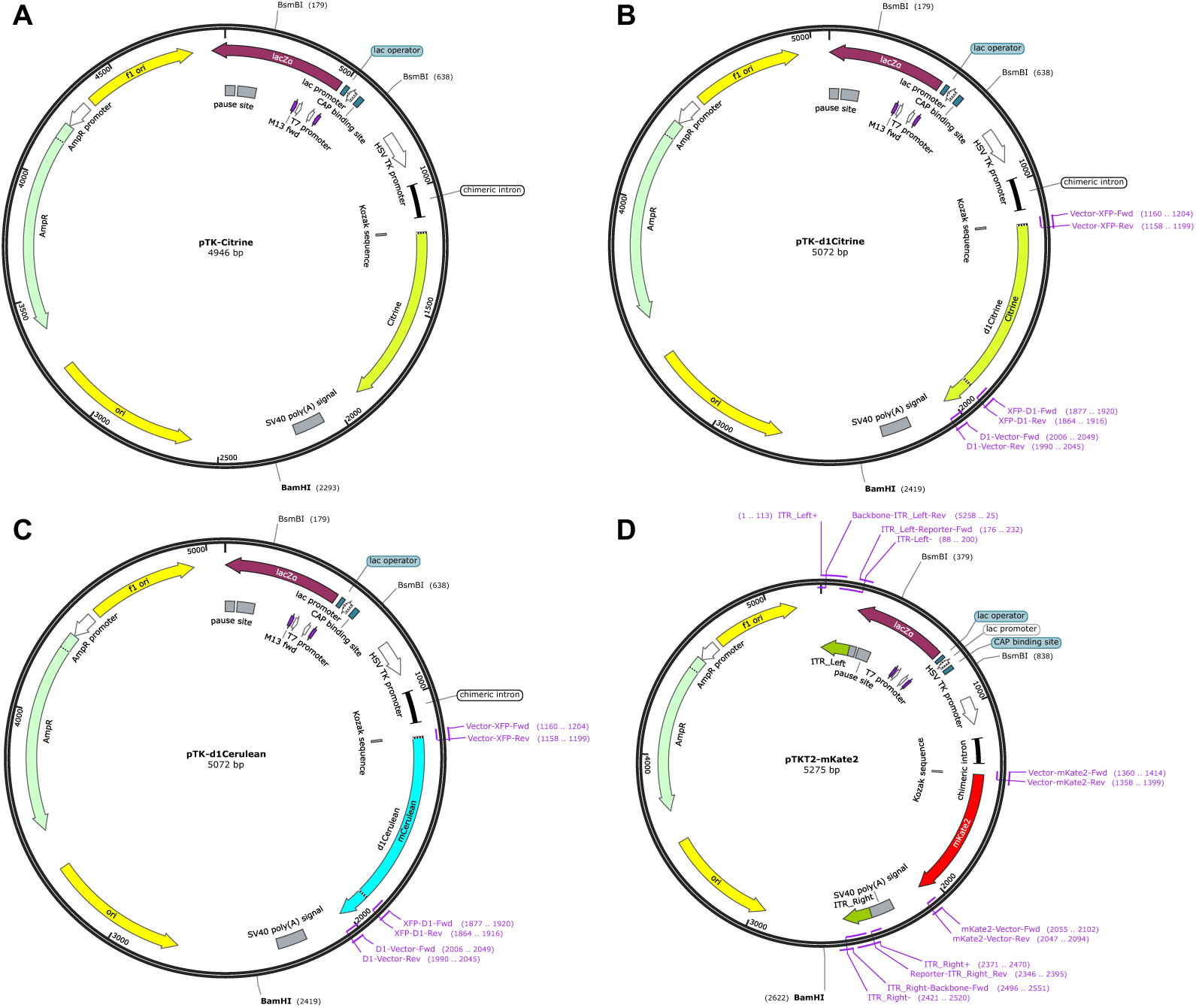
Maps of reporter plasmid vectors used to generate synthetic reporter constructs. **A)** pTK-Citrine. **B-D)** Vectors generated via Gibson Assembly. Each map indicates the PCR primers/oligos used. Sequences are listed in Supplementary Table 4. **B)** pTK-d1Citrine and **C)** pTK-d1Cerulean encode destabilised fluorescent proteins. **D)** pTKT2-mKate2 includes left and right ITRs for genomic integration of the reporter cassette via Tol2 transposase.

**Supplementary Figure 3:**
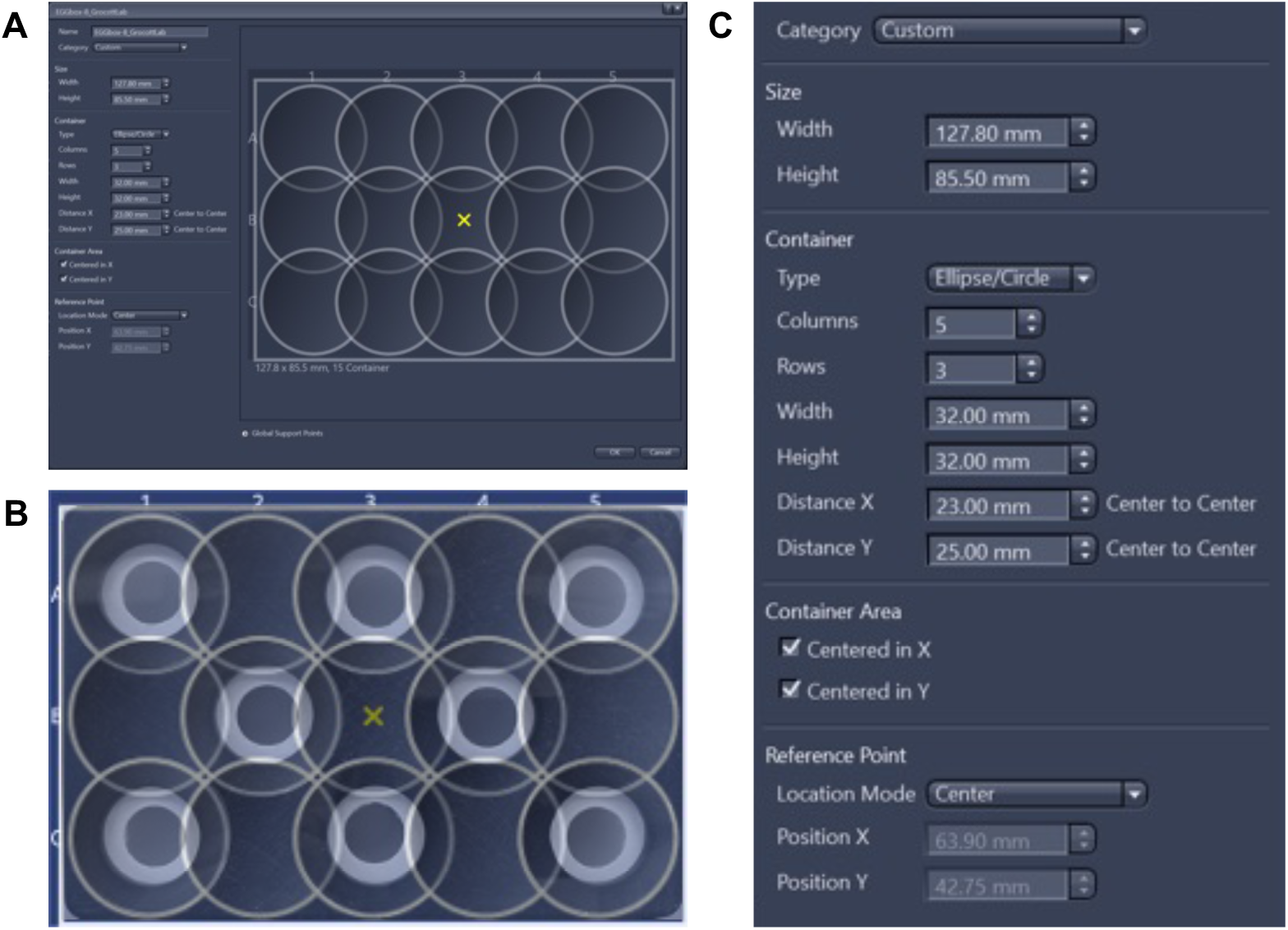
Custom Zen Blue sample carrier specification for use with EGGbox-8. Zen Blue does not provide for sample carriers with offset wells. Thus, we specified **A)** a custom sample carrier with a regular grid-pattern of overlapping wells. **B)** Overlay showing how top, middle, and bottom rows of EGGbox-8 respectively intersect with odd, even, and odd columns of the custom sample carrier. The reference point for stage calibration is the exact centre of the plate (yellow X). **C)** Dimensions of custom sample carrier.

**Supplementary Table 1:**
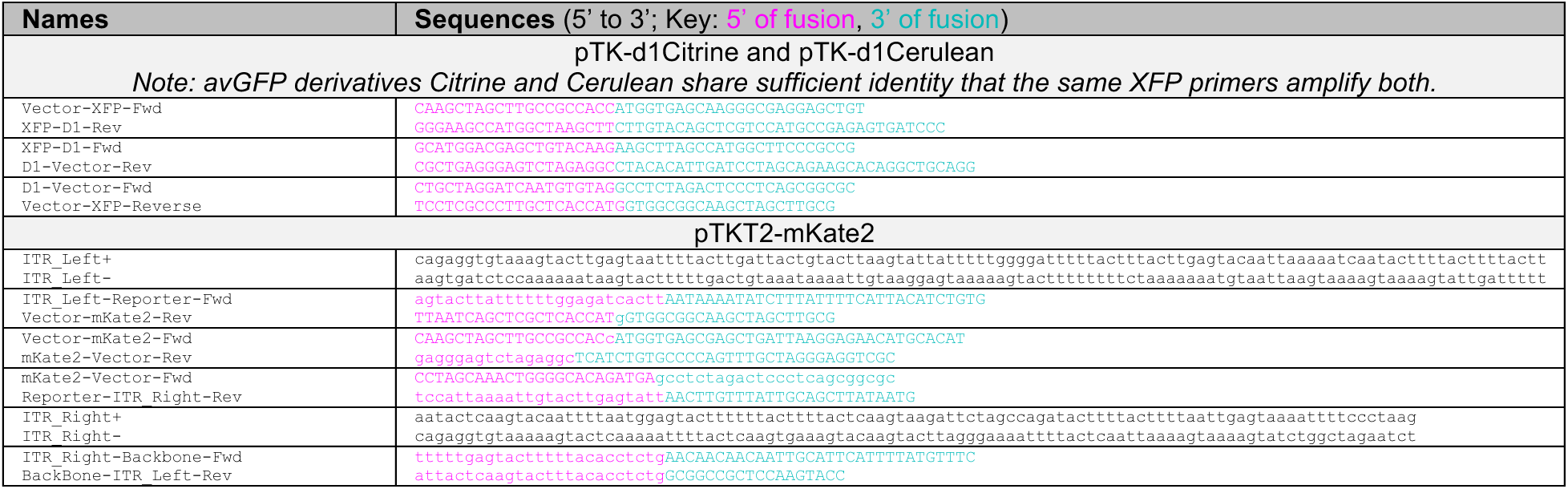
PCR primer and oligo pairs used for Gibson Assembly of novel reporter plasmid vectors.

**Supplementary Table 2:**
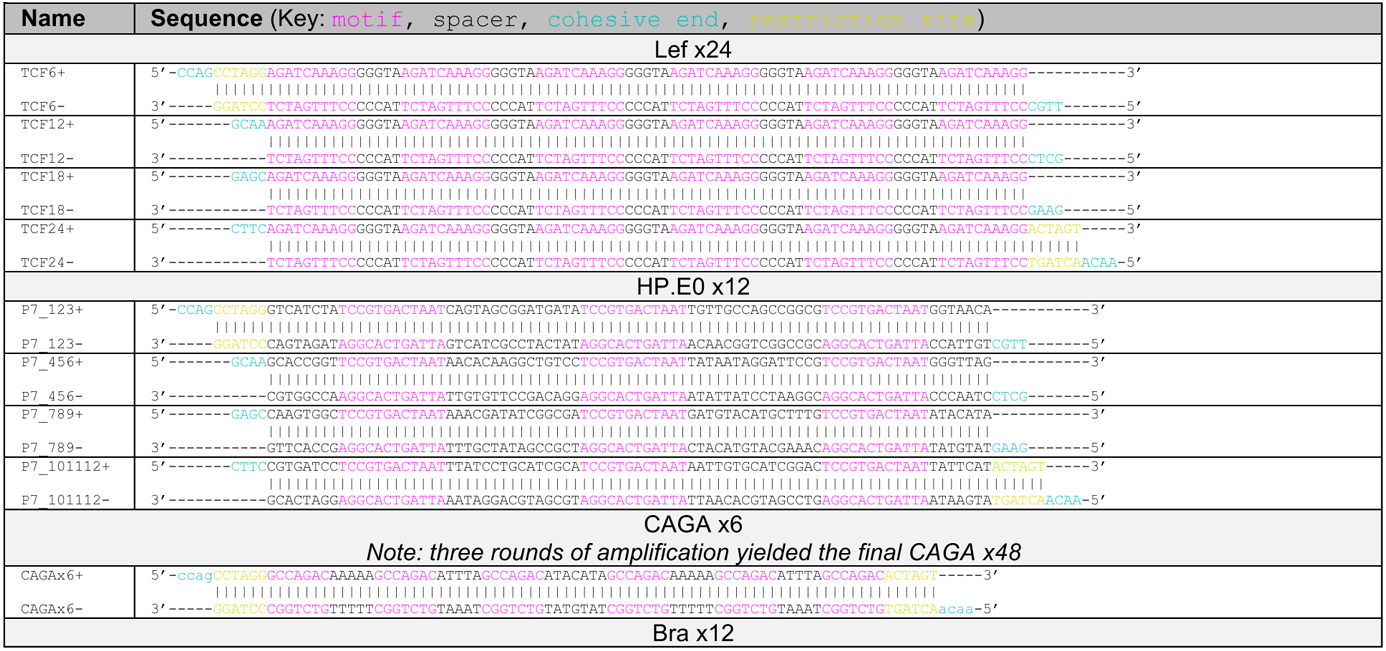
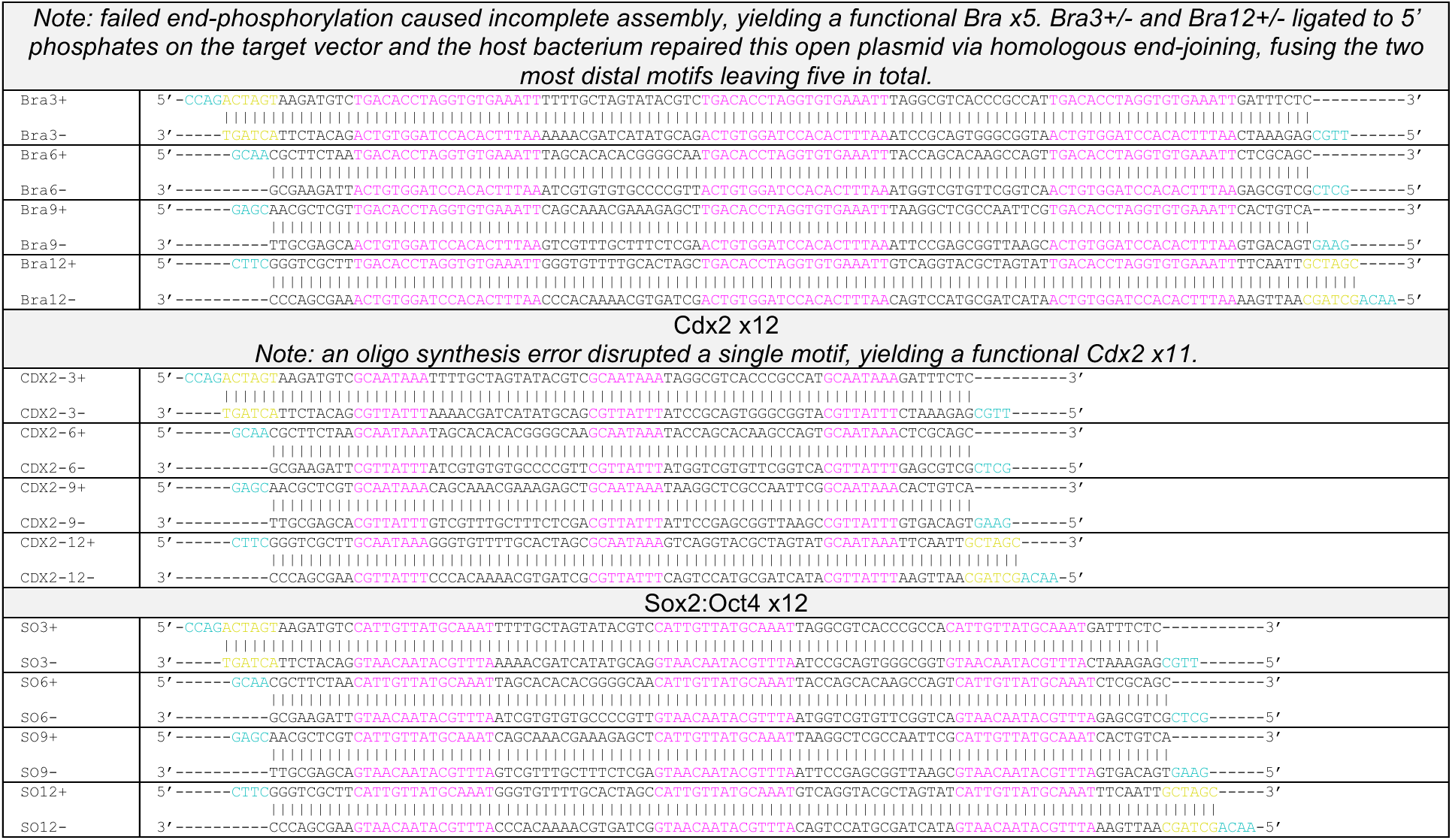
ssDNA oligos were end-phosphorylated and annealed for Golden Gate assembly of synthetic enhancers.

**Supplementary Table 3:**
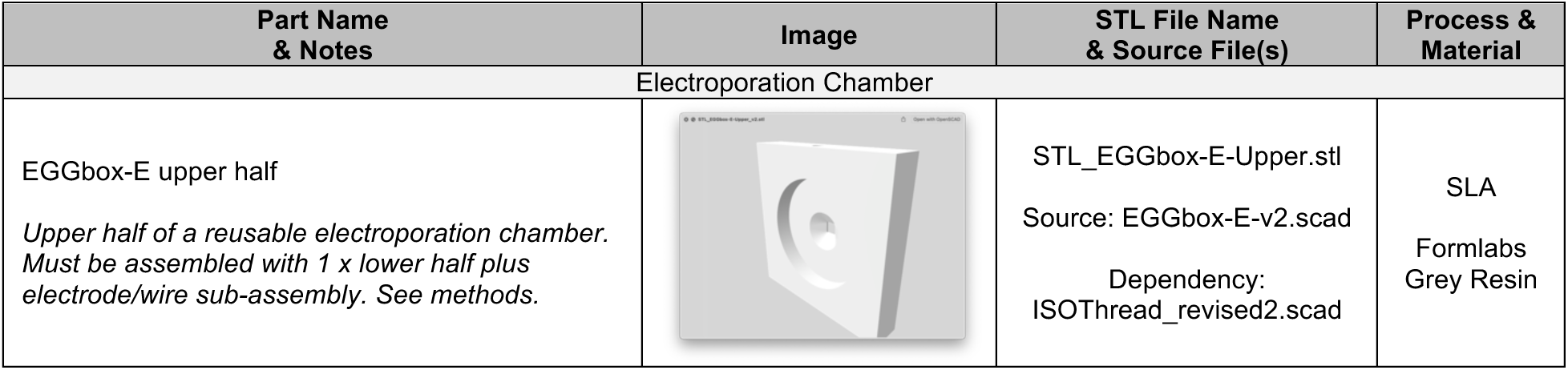
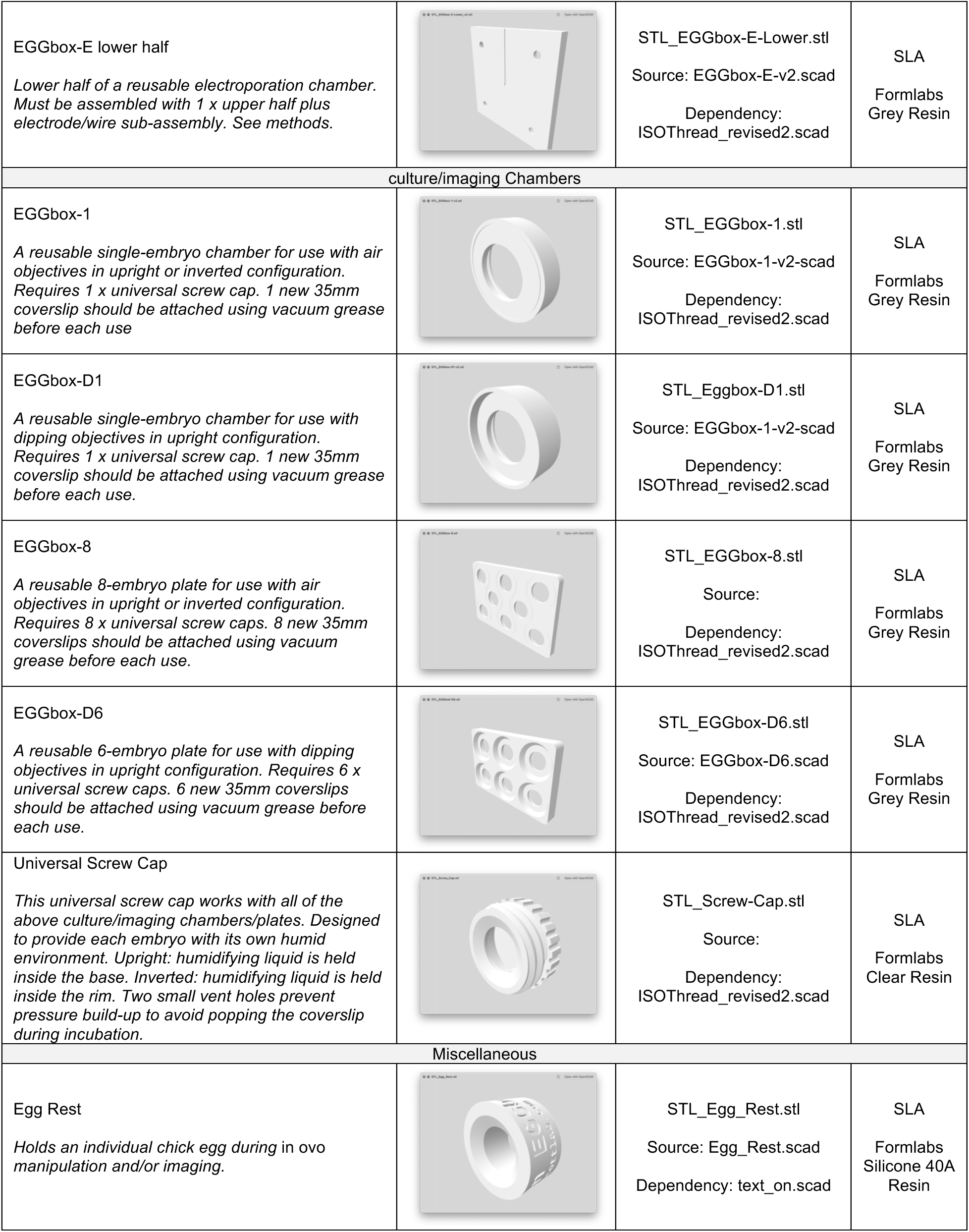
List of 3D-printable parts:

| Part Name & Notes | Image | STL File Name & Source File(s) | Process & Material |
| --- | --- | --- | --- |
| Electroporation Chamber |  |  |  |
| EGGbox-E upper half<br>*Upper half of a reusable electroporation chamber. Must be assembled with 1 x lower half plus electrode/wire sub-assembly. See methods.* |  | STL_EGGbox-E-Upper.stl<br>Source: EGGbox-E-v2.scad<br>Dependency: ISOThread_revised2.scad | SLA<br>Formlabs Grey Resin |
| EGGbox-E lower half<br>*Lower half of a reusable electroporation chamber. Must be assembled with 1 x upper half plus electrode/wire sub-assembly. See methods.* |  | STL_EGGbox-E-Lower.stl<br>Source: EGGbox-E-v2.scad<br>Dependency:<br>ISOThread_revised2.scad | SLA<br>Formlabs<br>Grey Resin |
| culture/imaging Chambers |  |  |  |
| EGGbox-1<br>*A reusable single-embryo chamber for use with air objectives in upright or inverted configuration. Requires 1 x universal screw cap. 1 new 35mm coverslip should be attached using vacuum grease before each use* |  | STL_EGGbox-1.stl<br>Source: EGGbox-1-v2.scad<br>Dependency:<br>ISOThread_revised2.scad | SLA<br>Formlabs<br>Grey Resin |
| EGGbox-D1<br>*A reusable single-embryo chamber for use with dipping objectives in upright configuration. Requires 1 x universal screw cap. 1 new 35mm coverslip should be attached using vacuum grease before each use.* |  | STL_Eggbox-D1.stl<br>Source: EGGbox-1-v2.scad<br>Dependency:<br>ISOThread_revised2.scad | SLA<br>Formlabs<br>Grey Resin |
| EGGbox-8<br>*A reusable 8-embryo plate for use with air objectives in upright or inverted configuration. Requires 8 x universal screw caps. 8 new 35mm coverslips should be attached using vacuum grease before each use.* |  | STL_EGGbox-8.stl<br>Source:<br>Dependency:<br>ISOThread_revised2.scad | SLA<br>Formlabs<br>Grey Resin |
| EGGbox-D6<br>*A reusable 6-embryo plate for use with dipping objectives in upright configuration. Requires 6 x universal screw caps. 6 new 35mm coverslips should be attached using vacuum grease before each use.* |  | STL_EGGbox-D6.stl<br>Source: EGGbox-D6.scad<br>Dependency:<br>ISOThread_revised2.scad | SLA<br>Formlabs<br>Grey Resin |
| Universal Screw Cap<br>*This universal screw cap works with all of the above culture/imaging chambers/plates. Designed to provide each embryo with its own humid environment. Upright: humidifying liquid is held inside the base. Inverted: humidifying liquid is held inside the rim. Two small vent holes prevent pressure build-up to avoid popping the coverslip during incubation.* |  | STL_Screw-Cap.stl<br>Source:<br>Dependency:<br>ISOThread_revised2.scad | SLA<br>Formlabs<br>Clear Resin |
| Miscellaneous |  |  |  |
| Egg Rest<br>*Holds an individual chick egg during in ovo manipulation and/or imaging.* |  | STL_Egg_Rest.stl<br>Source: Egg_Rest.scad<br>Dependency: text_on.scad | SLA<br>Formlabs<br>Silicone 40A<br>Resin |

**Supplementary Table 4:**
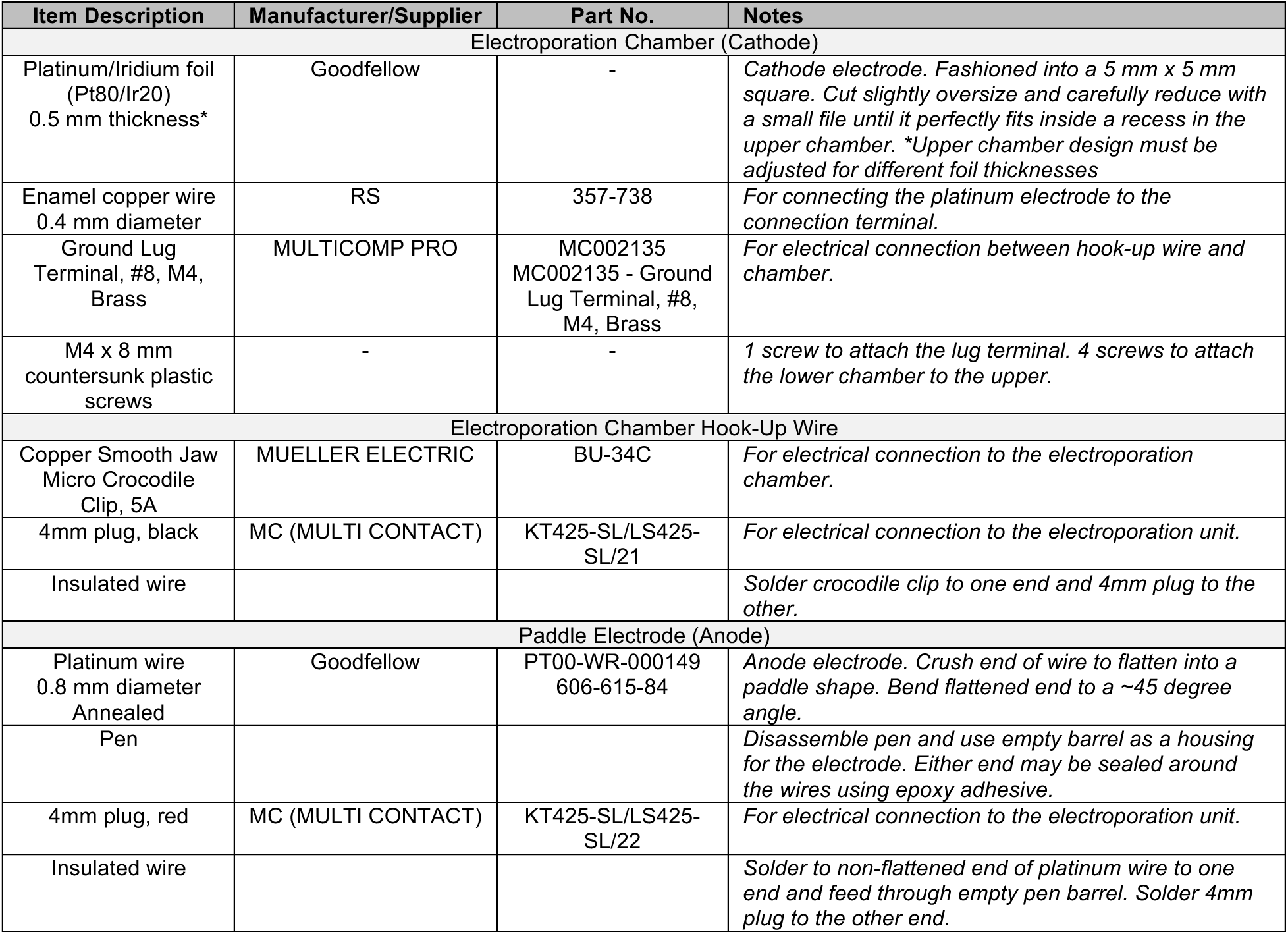
List of materials for assembling electroporation chamber, hook-up wire and paddle electrode.

**Supplementary Table 5:** List of materials required for assembling reusable culture/imaging chambers.

| Item Description | Manufacturer/Supplier | Part No. | Notes |
| --- | --- | --- | --- |
| Thickness No 1, 32 mm diameter round, borosilicate glass coverslips, 100 pieces | VWR | 631-0182 | Coverslips ensure greatest sample stability during imaging |
| High vacuum grease, 50g | Dow Corning | - | For attaching coverslips onto chambers/plates |
| Xtra-Clear Advanced Polyolefin StarSeal Film, 100 pieces | Starlab | E2796-9795 | Alternative to coverslip/grease. Self-adhesive low-autofluorescence film for microscopy. Less sample stability than coverslips (will drift more in z), but easier/quicker setup/clean-up. |
| Sodium Chloride (NaCl) | Fisher | BP358-212 | For making simple saline. Dissolve 7.19 g in 1 litre of sterile H<sub>2</sub>O and autoclave. |
| Agar (High Strength) | Melford |  | Heat 0.72 g in 120 ml of simple saline until dissolved and cool to 48 degC in water bath. |
| Thin albumin collected fresh from eggs, or stored at -20 degC. | - | - | Warm to 48 degC and mix 1:1 with 0.6% agar to form albumin-agar solution. |
| Penicillin/Streptomycin (100x) | Fisher | 15140-122 | Dilute 1/100 into albumin-agar solution. |
| Ringer's solution tablets | Fisher | R/0350/79 | For collecting and cleaning embryos. Dissolve 1 tablet in 500 ml of sterile H<sub>2</sub>O. |
| Whatman 3MM CHR chromatography paper | Whatman | 3030-909 | For harvesting chick embryos using the EC method. |
| 0.5-inch hole punch | EK Success | 54-10062 | For fabricating filter paper rings with 1/2" inner diameter. |
| 0.75-inch hole punch | Efco | 1791114 | For fabricating filter paper rings with 3/4" inner or outer diameter. |
| 1.5-inch hole punch | Efco | 1793010 | For fabricating filter paper rings with 1 1/2" outer diameter. |

